# ShapeCluster: Applying parametric regression to analyse time-series gene expression data

**DOI:** 10.1101/035782

**Authors:** Philip Law, Vicky Buchanan-Wollaston, Andrew Mead

## Abstract

High-throughput technologies have made it possible to perform genome-scale analyses to investigate a variety of research areas. From these analyses, vast amounts of data are generated. However, this data can be noisy, which could obscure the underlying signal. Here, a high-throughput regression analysis approach was developed, where a variety of linear and nonlinear parametric models were fitted to gene expression profiles from time course experiments. These models include the logistic, Gompertz, exponential, critical exponential, linear+exponential, Gaussian and linear functions. The fitted parameters from these models reflect aspects of the model shape, and thus allowed for the interpretation of gene expression profiles in terms of the underlying biology, such as the time of initial gene expression. This provides a potentially more mechanistic approach to studying the genetic responses to stimuli. Together with a cluster analysis, termed ShapeCluster, it was possible to group genes based on these aspects of the expression profiles. By investigating different combinations of parameters, this added flexibility to the analysis and allowed for the investigation of the data in multiple ways, including the identification of groups of genes that may be co-regulated, or participate in response to the biological stress in question. Clusters from these methods were assessed for significance through the use of over-represented annotation terms and motifs, and found to produce biologically relevant sets of genes. The ShapeCluster package is available from https://sourceforge.net/projects/shapecluster/.

## 2 INTRODUCTION

Time series experiments are a popular means of investigating the dynamic changes in an organism's transcriptome in response to some stimulus. This type of data may be cross-sectional or longitudinal. Cross-sectional data indicates that each data point was obtained from an independent sample (e.g. leaf samples at different times), whereas in longitudinal data, subsequent data points are obtained from the same individual (e.g. blood samples for a patient at different times). In time series data, there is an obvious dependence of each observation on the past observations. Numerous techniques have been developed to take this temporal information into account, including the use of Bayesian-based hierarchical clustering algorithms (e.g. Cooke, et al. (2011) and Heard, et al. (2006)), hidden Markov model (HMM) algorithms (e.g. Oh, et al. (2013) and Schliep, et al. (2003), and curve fitting using smoothing spline clustering models (e.g. Déjean, et al. (2007) and Ma, et al. (2006)). These techniques are able to model any unknown shape with a relative amount of ease, without requiring any prior information about the structure of the data. However, these methods may not have a biologically relevant interpretation.

Parametric regression analysis is a common technique that has been applied to multiple fields of science, including ecology (e.g. Dalbiés-Dulout and Doré (2001) and Schoolfield, et al. (1981)), analytical chemistry (e.g. Watkins and Venables (2006)), and medical statistics (e.g. Woolcock, et al. (1984)) where a specific model is fitted to some data. In all these cases, a parametric model was used to describe the relationship between the response and the predictor.

Regression analyses can be used to obtain a better explanation into the function and mode of operation of genes, by using the parameters to provide insights into differences between sets of genes, or indicating when particular events occur (Eastwood, et al., 2008; Ma, et al., 2006; Watkins and Venables, 2006). For example, Watkins and Venables (2006) used the fitted values for a parametric model to identify the optimal separation point and pH of four related carboxylic acids. Similarly, Eastwood et al. (2008) used the critical-exponential model to describe the expression changes in a number of genes, and used the fitted parameters to identify the time of maximal transcript level, and identify the asymptotic response level.

Here an approach is described where the fitted parameters from a regression analysis are used to infer biological interpretation directly from the shape of the expression profiles. Using the parameters in a cluster analysis provides a means of analysing sets of genes on aspects of the shape, thus providing a more mechanistic approach to study the genetic responses to stimuli.

## 3 METHODS

Data was obtained from a high-resolution time-series in Arabidopsis investigating the molecular effects of senescence (Breeze, et al., 2011). The gene expression data was fitted to a number of model shapes, namely logistic, Gompertz, exponential, critical exponential, linear+exponential, and Gaussian and linear. The models were fitted using the *nls* and *Im* functions in the R statistics package, together with the use of self-starter functions. The self-starters were developed using aspects of the curves to identify approximate estimates of parameter values, and is based on the concept of stable parameters (Ross, 1970).

To determine the quality of the fitted curve, goodness-of-fit statistics were used, primarily the lack of fit sum of squares (SS_lack-of-fit_)· For any regression model, it is possible to partition the total sum of squares into the regression sum of squares (SS_regression_) and the residual sum of squares (SS_residual_). The SS_residual_ can then in turn be partitioned into the pure error sum of squares (SS_pure error_) and the SS_lack-of-fit_.

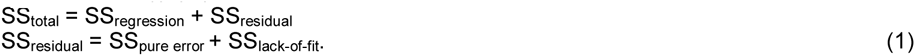

The SS_lack-of-fit_ value was calculated by fitting a saturated model to the time-series data, so named as a parameter is allowed for each time point, and is thus saturated with parameters. This model describes the expected response at each time point, and represents a model that is formulated with no assumptions with regards to the response shape and time dependence of the data points. Like SS_total_, the saturated model total sum of squares can be decomposed into two parts. However, in this case, the residual sum of squares is equal to the SS_pure error_. By using these values, together with equation (1), the SS_lack-of-fit_ for the regression model can be calculated

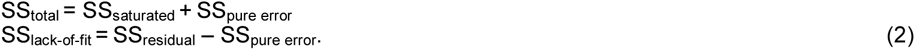

An R^2^ value (coefficient of determination) was also calculated to evaluate goodness-of-fit by comparing the residual sum of squares to the total sum of squares. Similarly, using the SS_lack-of-fit_ it is possible to calculate an 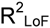 value based on the lack-of-fit. Thus

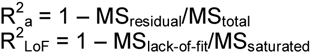

where MS_residual,_ MS_total_ MS_lack-of-fit,_ MS_saturated_ are the residual mean square, total mean square, lack-of-fit mean square and saturated mean square, respectively.

In addition, an F-test was used to determine how appropriate the model is given the data.

All the results of the regression analysis, including gene information, best fit information and fit statistics were stored in an SQLite database. These results and data could then be accessed for further analysis, including the cluster analysis. For this analysis, a distance matrix was created using the fitted parameters. The standard error of estimation for the parameters was used to filter the fits. The clusters were determined using the *hclust* function in R, using an average linkage distance. Clusters were identified using the Dynamic Hybrid algorithm from the Dynamic Tree Cut package (Langfelder, et al., 2008).

To cluster using multiple parameters, a number of different methods were used. The simplest case is “simultaneous clustering”, where a Euclidean distance was used to create a single combined distance matrix of the parameters of interest. In addition, two meta-clustering approaches were used, namely sequential and cross meta-clustering. In “sequential” metaclustering, the shapes are clustered first on a single parameter. Following this, the members of each of the clusters are further clustered based on a second parameter. With “cross” clustering, the clusters were formed using each of the two parameters independently, thus resulting in two sets of clusters. The genes contained within each cluster in one set were compared to the genes contained in every other cluster in the other set, in a pair-wise manner (i.e. cross classification of the cluster memberships). If a gene is contained in a pair of comparisons, the gene is assigned to this intersection of two clusters. Clusters were evaluated for biological significance using the *goStats* package (Falcon and Gentleman, 2007), and the BHI function in the *clValid* package (Brock, et al., 2008). Metabolic pathway analysis was performed using the mappings of gene products to metabolic pathways in MAPMAN (Thimm, et al., 2004).

The presence of over-represented upstream regulatory sequences was determined using the promoter analysis as described in Breeze et al. (2011). In brief, 1,037 binding motifs in plants were obtained from the JASPAR (Mathelier, et al., 2014) and PLACE (Higo, et al., 1999) databases. Motifs were identified from sequences 500 bp upstream of the transcription start site of each gene in the cluster. For each motif, the frequency in the cluster was computed, and compared to the frequency in the Arabidopsis genome. A hypergeometric test was used to provide a description of the significance of the presence of a motif.

## 4 RESULTS

The above regression models were fitted to data from a high-resolution time-series microarray experiment in Arabidopsis investigating the molecular effects of long day senescence (Breeze, et al., 2011). In brief, the data consisted of 11 time points, and 4 replicates at each time point. The microarrays contained over 32 500 probes, which mapped to approximately 24 000 unique gene models.

### 4.1 Fitting parametric models to gene expression profiles

Using a least-squares approach, eight different models with distinct shapes, namely logistic, Gompertz, exponential, critical exponential, linear+exponential, and Gaussian and linear models were fitted to the senescence gene expression profiles. The best fitting model was determined using the AIC. Following (Burnham and Anderson, 2002), any models with a difference in AIC values (ΔΑIC) less than 2 to the smallest AIC value were retained, meaning an expression profile can fit to multiple models.

The parameters from these models influence aspects of the shape of the curve, which can be used to interpret functional aspects of the gene expression profiles. For example, the logistic function represents a symmetric sigmoid curve characterised by a rapid growth rate in the beginning, slowing down to a constant growth rate, before finally approaching the asymptotic maximum value. The *a* parameter describes the starting expression level, *b* represents the distance between asymptotes (range of gene expression response), *m* is the time (*t*-value) at which maximum growth is reached, and *s* is related to the slope at time *m*. Fig 1 shows an example of an expression profile from the dataset fitted by the logistic model, together with an indication of the aspects of the curve that are affected by the parameter values. A decreasing response is also possible with a maximum initial value followed by rapid decrease to a minimum asymptote, and is determined by the signs of the fitted parameter values. If the *s* and *b* parameters have the same sign, the curve is increasing, whereas if they are of opposite signs, it represents a decreasing response.

**Fig. 1.**
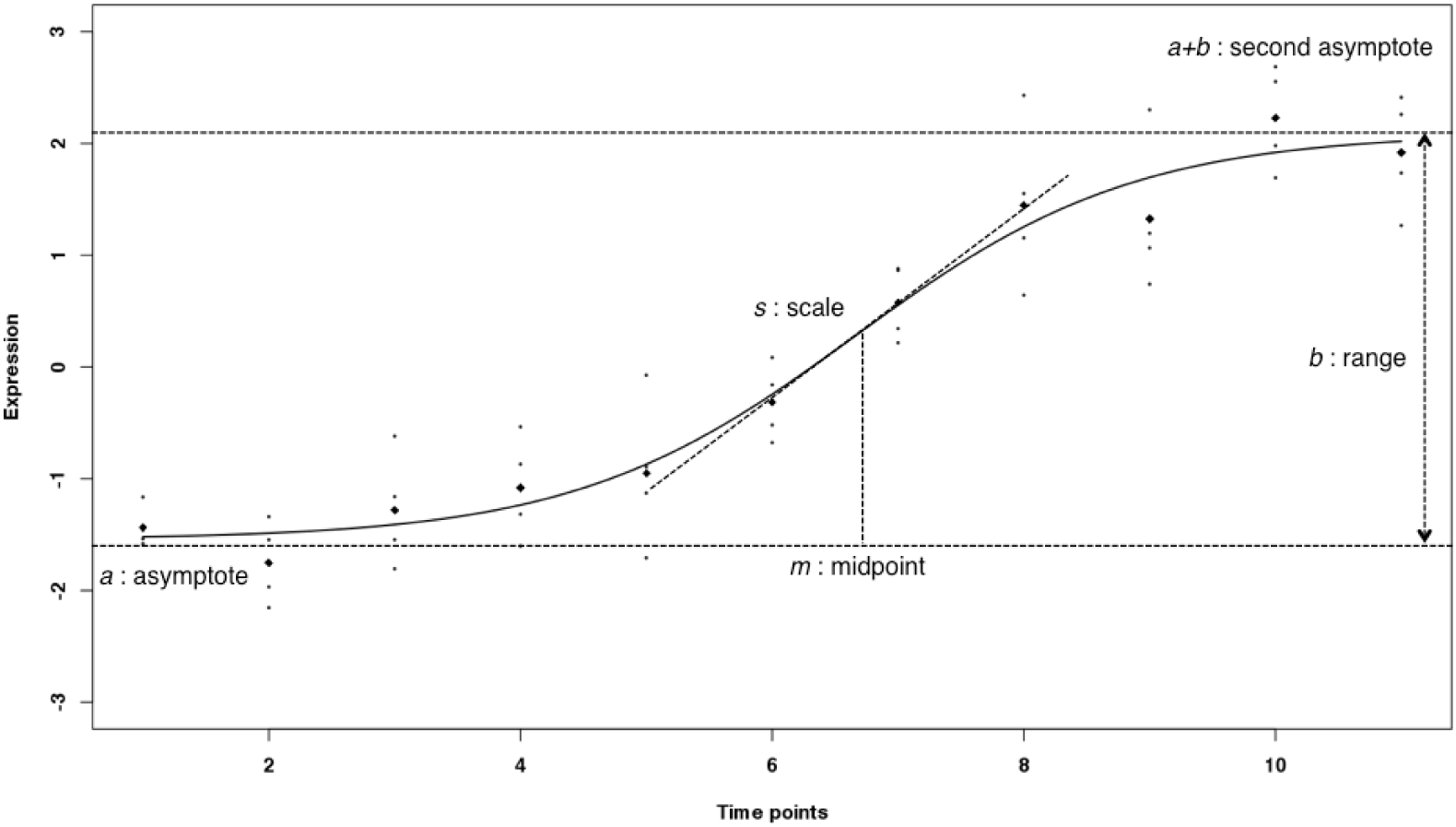
Example of a logistic curve (*y = a + b/(1+exp((m-t)/s)))* fitted to a gene expression profile with the parameters illustrating the aspects of the curve they influence. The small dots are the replicate samples and the black diamonds are the means of each time point.

While the parameters from some models are directly informative, it may be desirable to calculate other parameter estimates that describe alternative aspects of the curves. For example, for the sigmoid models, the values of the function increase (or decrease) from one asymptotic value to another. It is thus possible to find the time point (*t*-value) when 5% of the difference between asymptotes has been reached (denoted the *5per* parameter), indicating the point at which the increase (or decrease) is starting. In terms of gene expression, this may indicate the point at which the gene is being activated or repressed. To calculate the *5per* parameter, this would be the point where *y = a +0.05*b*. Thus, using the equation for the logistic function (Fig. 1), the time point where this value occurs is *5per = m − s*log(1/0.05 − 1)*. Similarly other parameters such as the *grad* parameter to describe the rate of change of gene expression at time *m* can be defined.

The goodness-of-fit was also calculated using the adjusted R^2^ (R^2^_a_), lack-of-fit R^2^ (R^2^_LoF_), and the F-test p-value, and could be used to filter out those fitted profiles with a poor fit to the models. The standard error of estimation was also used to filtered parameters that were poorly estimated. For the derived parameters, the delta method was used to estimate the standard error (Fox, 2002; Fox, 2008; Ritz and Streibig, 2008).

### 4.2 Cluster results

The parameters from the fitted models were further used to group those genes that had a similar aspect in their expression profiles. For example, identifying genes that become activated at the same time, or change at the rate. This analysis was named ShapeCluster.

In contrast to conventional cluster analyses, ShapeCluster examines the fitted curves rather than the observed data. The algorithm operates in a two-step process: first, sets of genes are identified based on the particular model that fitted the gene expression profiles; and second, the similarity of genes based on one or more of the biologically interpretable parameters is determined. Several methods of producing clusters with combinations of parameters were developed, namely simultaneous parameter clustering, sequential meta-clustering, and cross meta-clustering. The simultaneous parameter calculation was performed by combining the parameter values using a Euclidean distance to produce a single measure. The sequential meta-cluster analysis clustered on the first parameter, and then these clusters are subsequently clustered based on a second parameter. The cross meta-cluster was performed by independently clustering the genes on each of the two parameters, and the genes in common between the two clusters are identified.

In order to only perform the cluster analysis on fits that had good fits, the goodness-of-fit statistics were used to filter the fits. Thus, thresholds of R^2^_LoF_ > 0.6, R^2^_a_ > 0.6, and F-test p-value < 0.05 were used. These values were determined to be stringent enough to remove inadequate fits, but still flexible enough to allow a sufficient proportion of the genes through. However, these thresholds are arbitrary and can be adjusted accordingly.

Significant results from the analysis of the senescence data, using the above thresholds are summarised below. Described are the results from the logistic, exponential, and Gaussian shapes.

For the genes that fitted the logistic shape, the profiles were clustered using the *5per* parameter (time of 5% of maximum response), which provided insights into genes that were up- or down-regulated at a given time, and using this parameter made it possible to determine when specific sets of genes were activated or repressed. This in turn provided a means of identifying what biological processes were being activated or repressed in response to the stimulus, allowing the times that metabolic functions occurred to be elucidated. In contrast, performing the cluster analysis using the *grad* parameter would identify genes that have gene expression changing at the same rate, and thus may possibly be co-regulated by a common regulator or transcription factor. Fig 2 shows an illustration of a cluster formed using these two parameters.

**Fig. 2.**
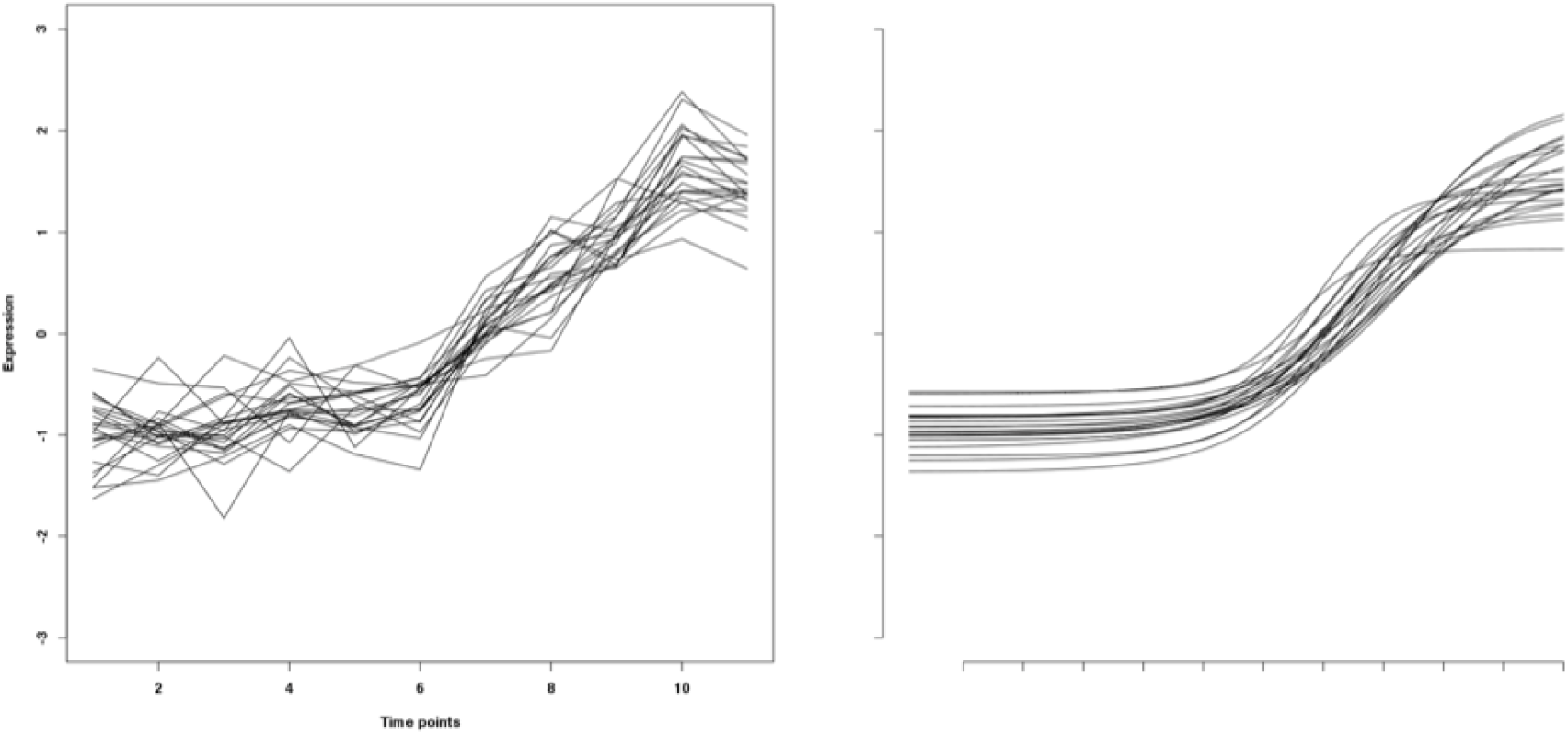
A representation of the clusters obtained clustering the logistic curves, using the *5per* and *grad* parameters. The figure on the left was plotted using the mean raw expression data, and the figure on the right shows the smoothed curves using the fitted parameters.

Investigating over-represented GO terms found that the genes with increasing responses and a small *5per* parameter value (genes with the early activation times) were involved with stress responses and nucleic acid metabolism, whereas genes with later activation times were involved in the ageing response. Interestingly, in the latter clusters, there were genes involved in auxin homoeostasis, such as AUXIN UPREGULATED 3 (AUR3, AT4G37390). Auxins are a group of plant hormones, which are involved in a number of developmental processes in plants, including senescence (Osborne, 1959), so this set of genes may be involved in regulating the senescence responses. The genes with decreasing responses were related to photosynthesis and chloroplasts, with several clusters showing different times of down-regulation and rates of change, indicating multiple stages of down-regulation of photosynthesis.

In the genes that fitted the exponential shape, the profiles were clustered using the rate of change parameter (*r*) and concavity (*b*) parameters. The *r* parameter was selected as it would identify genes that have a similar rate of change in their gene expression, and thus may be involved in the same biological processes. The *b* parameter determines if the exponential shape will be concave or convex, indicating that the gene expression level is increasing to a plateau (*b*<0), or decreasing to the asymptote (*b*>0). Analysing the annotation terms revealed that the genes with a concave increasing shape were enriched in terms involving transporter activity. Conversely, the genes with a convex decreasing shape were enriched for ribosomes and RNA processing, suggesting that as the plant ages, the ribosomal and translational activities decrease. In addition, in clusters with lower *r* values (more linear response), the genes with an increasing profile were enriched for terms relating to stress responses and water deprivation, and could be involved in the activation of senescence responses, while those with a decreasing response were involved in photosynthesis.

The Gaussian model describes a change in expression to a maximum, or minimum, followed by a transition back to the starting expression level. Here, the *m* parameter provides an indication of the time when an up-regulated gene becomes down-regulated, or *vice versa*, whereas the s parameter provides an indication of the duration of the gene expression response. Thus the *m&s* clusters would identify genes that have a maximum (or minimum) at the same time, as well as having a similar spread in the data points. This reflects a transient response that is being activated or repressed in response to some biological signal.

In the expression profiles that increase to a maximum, the clusters were enriched for terms such as carbohydrate metabolism, transporter activity, and vacuole regulation and metabolic activities. This activity indicates that the plant is beginning to activate the transport processes in order to mobilise the macronutrients to other parts of the plant, such as storage organs. Notably, genes in clusters with a large *m* parameter (late maximum response time) were enriched for transcription factor (TF) activity, indicating TFs that are activated near the end of the senescence process. These clusters contained the genes *ANAC014 (AT1G33060)* and *ANAC089 (AT5G22290)*. The NAC TF family has been shown to be involved in the senescence process (Breeze, et al., 2011; Hickman, et al., 2013), so it is possible that these genes are involved in the regulation of the senescence response. ANAC089 is involved in regulating the flowering time in Arabidopsis (Li, et al., 2010), and ANAC014 currently has no known biological function. In contrast, in the genes decreasing to a minimum, the clusters contained genes that were involved with RNA binding activity, and again being involved with photosynthesis and chloroplasts. The former set of clusters were all down-regulated early in the time series (around time point 2–4), and were primarily involved in translation. The latter clusters were repressed at a much later stage (after time point 10) indicating that the photosynthesis genes are becoming down-regulated towards the end of the time series, as discussed above.

In addition to the simultaneous parameter clustering, the meta-clustering approaches were also applied to the expression profiles. In general, the over-represented annotation terms were the same as the simultaneous clustering, described above. A few notable terms included those related to the response to abscisic acid (ABA) in the Gaussian models, using the sequential meta-clustering. These genes had an increasing response, reaching a maximum at around time point 10 before becoming down-regulated again. This is consistent with other findings, where it has been shown that there is an accumulation of ABA due to the up-regulation of ABA biosynthetic genes during senescence (Breeze, et al., 2011; Buchanan-Wollaston, et al., 2005; van der Graaff, et al., 2006). Another term that was not seen with the simultaneous parameter clustering is the presence of genes that were involved in ethylene mediated signalling pathways. These genes were again found using the Gaussian models, and sequential meta-clustering. The genes initially decreased until around time point 2 before becoming up-regulated. It has been shown that ethylene levels increase during the senescence process, due to the up-regulation of ET biosynthetic genes as the plant ages (van der Graaff, et al., 2006).

Using the average parameter values from each cluster, as well as the information described above, a simple time line of the biological processes that were occurring over time can be inferred during the senescence process, by using the timing parameters from the different shapes (e.g. *5per* and *m*). By using these timing parameters together with the over-represented annotation terms, it was possible to determine when specific biological events were taking place. These results were similar to those found in the published results from the senescence time course (Breeze, et al., 2011). In particular, responses to water deprivation and pectinesterases were up-regulated at the same points. Down-regulated in both were genes involved in amino and nucleic acid metabolism, as well as several series of photosynthesis related genes. There were however a few differences. While both analyses identified chlorophyll related genes being down-regulated between time points 5–7, ShapeCluster did not identify photosynthesis related genes that were down-regulated at time point 3. It did however identify photosynthesis genes that are down-regulated later, at time point 9. Other new discoveries include the identification of early up-regulation of ethylene signalling, auxin homeostasis at time point 7, and late ABA signalling.

Many methods to determine the accuracy of the clustering exist, such as the Rand Index and correlation based methods, where it is assumed that clusters are formed based on the entire profile. Here, Biological Homogeneity Index (BHI) was used (Datta and Datta, 2006). This algorithm attempts to evaluate a set of clusters based on how similar the annotations within each cluster are. The BHI scores for a variety of parameter and shape combinations from ShapeCluster are shown in Table 1, with an average BHI score of 0.37.

**Table 1.** Table of all the BHI scores for the cluster analyses performed on the senescence data, using different combinations of shape, direction, and parameter.

|  | Exponential | Gaussian | Gompertz | Logistic |
| --- | --- | --- | --- | --- |
| Single | *b*<br>0.35 | *m*<br>Inc: 0.34<br>Dec: 0.46 | *5per*<br>Inc: 0.37<br>Dec: 0.35 | *5per*<br>Inc: 0.35<br>Dec: 0.35 |
|  | *r*<br>0.34 | *s*<br>Inc: 0.33<br>Dec: 0.39 | *grad*<br>Inc: 0.37<br>Dec: 0.36 | *Grad*<br>Inc: 0.38<br>Dec: 0.37 |
| Simultaneous | *r&b*<br>0.34 | *m&s*<br>Inc: 0.37<br>Dec: 0.43 | *5per&grad*<br>Inc: 0.37<br>Dec: 0.39 | *5per&grad*<br>Inc: 0.37<br>Dec: 0.37 |
| Meta: cross | *r&b*<br>0.36 | *m&s*<br>Inc: 0.34<br>Dec: 0.42 | *5per&grad*<br>Inc: 0.36<br>Dec: 0.38 | *5per&grad*<br>Inc: 0.35<br>Dec: 0.40 |
| Meta: sequential | *r&b*<br>0.37 | *m&s*<br>Inc: 0.38 | *5per&grad*<br>Inc: 0.37 | *5per&grad*<br>Inc: 0.36 |

As a comparison, SplineCluster (Heard, et al., 2006), a Bayesian model-based hierarchical clustering algorithm was also used to cluster the data. To obtain as fair comparison as possible, the full set of 23,802 genes was filtered using the same R^2^_a_, R^2^_LoF_, and F-test p-value thresholds as before, resulting in a set of 8,216 genes. Application of SplineCluster to these genes, using the default parameters, resulted in a set of 98 clusters with a BHI score of 0.28.

### 4.3 Investigating co-clustering genes

As a further analysis of the cluster analysis, the clusters were examined to identify genes that co-cluster with a gene of interest. This gene was *PHOTOSYSTEM I LIGHT HARVESTING COMPLEX GENE 6 (LHCA6, AT1G19150)*, which encodes a light-harvesting complex I protein that forms part of photosystem I (Peng and Shikanai, 2011). It was thus expected that genes that cluster with it would also form part of the photosynthetic machinery.

The expression profile of *LHCA6* fitted to the decreasing logistic model, and was clustered using the *5per, grad*, and *5per&grad* parameters. By using these parameters, it was possible to identify genes that were repressed at the same time as *LHCA6* (using the *5per* parameter), genes that had the same rate of change in gene expression (*grad* parameter), and the genes that were being repressed at the same time as well as having the same rate of change *(5per&grad)*.

Genes that co-clustered using the *5per* parameter included a large number of photosystem I related genes, such as *PHOTOSYSTEM I SUBUNIT K (PSAK, AT1G30380)*, and *PHOTOSYSTEM I LIGHT HARVESTING COMPLEX GENE 5 (LHCA5, AT1G45474)*.

In the *grad* clusters, the genes *PHOTOSYSTEM II LIGHT HARVESTING COMPLEX GENE 2.3 (LHCB2, AT3G27690)* and *PHOTOSYSTEM II SUBUNIT T (PSBTN, AT3G21055)* were found, suggesting that components from both photosystems decrease at the same rate. In addition, genes encoding proteins involved in secondary metabolism were co-clustered, suggesting that multiple photosystem components are being regressed at the same rate as various metabolic processes. The multiple parameter clustering was performed using the simultaneous parameter clustering. Genes that co-clustered again included photosynthesis related genes, such as chlorophyll A-B-binding family protein (AT1G44575), as well as genes encoding receptor proteins.

As expected, the significant GO terms in all these clusters were related to the chloroplast (e.g. thylakoid, stroma, photosynthesis). In addition, it was found that the clusters contained genes that are involved in the photosystem I and II metabolic pathways.

The presence of common transcription factor binding sites in the upstream regions of these genes was also investigated (Fig 3). Transcription factor binding motifs in the upstream regions that were over-represented identified using a hypergeometric test, and having p<0.05. In the cluster grouped using the *5per* parameter showed that the MA0097.1 (TGACGTG) motif was over-represented. This motif is the binding site for the bZIP911 transcription factor. In the cluster grouped using the *grad* parameter, over-represented motifs included S-000503 (TCCAACCA), a UV-B responsive transcription factor; and S-000287 (TCCACGTGTC), the binding site of bZIP transcription factors which interact with an auxin-responsive gene, and in the cluster grouped using the both the *5per* and *grad* parameters, the binding site of a NFYA transcription factor subunit (MA0435.1, CCAAT). This family of transcription factors has been shown to be involved in the senescence process.

**Fig. 3.**
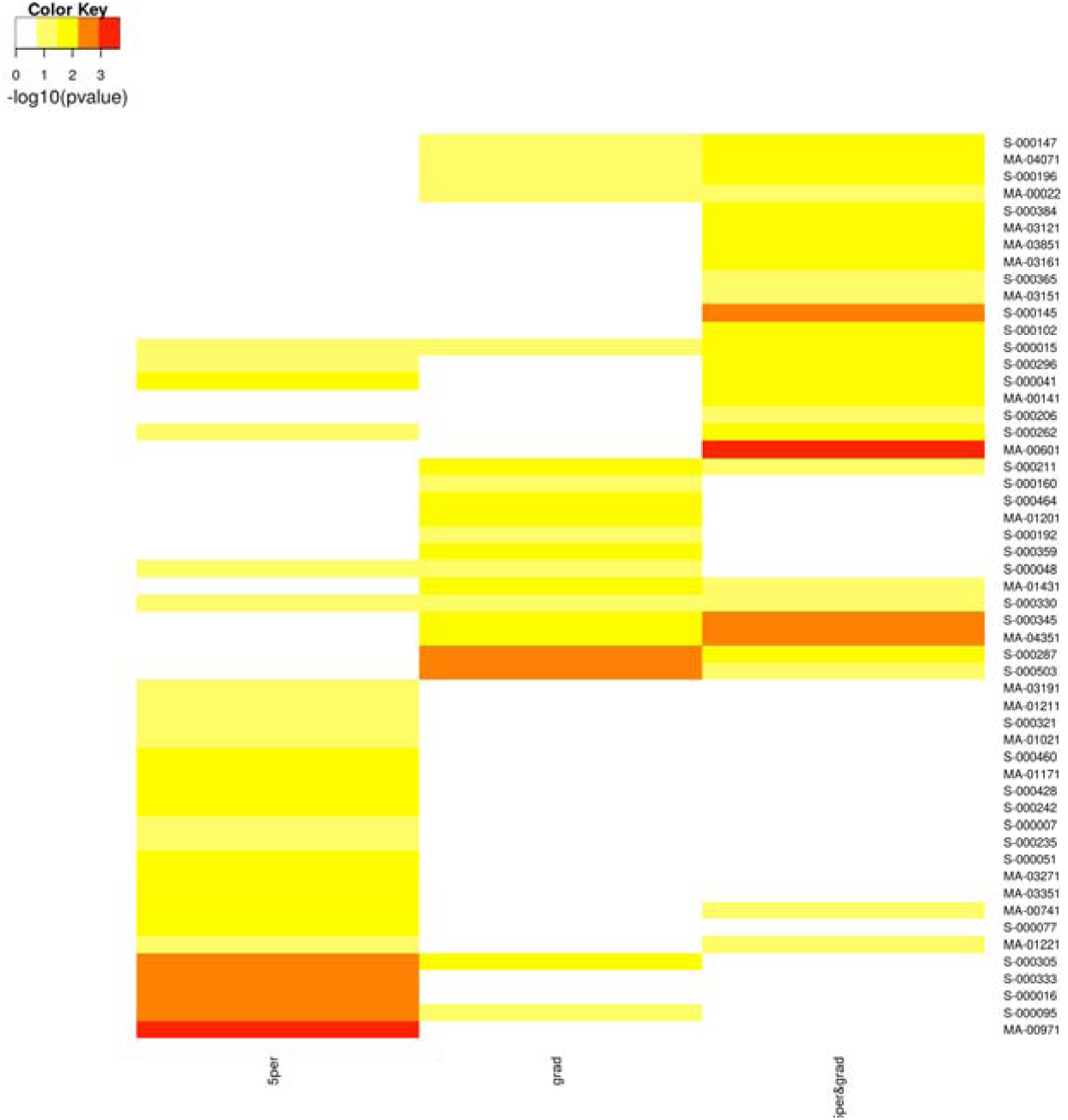
Heatmap showing the over-represented motifs in the sets of genes that co-cluster with *LHCA6* using the different parameters, namely *5per* (left), *5per&grad* (middle), and *grad* (right). Shown are the – log10(p-values) from the hypergeometric test.

## 5 DISCUSSION

Most clustering methods group observations together based on the profiles appearing the same. However, in a biological system, these methods may not identify genes that regulate each other. Kiddle, et al. (2010) presented a method of clustering time series expression data using an affinity propagation algorithm to identify profiles with a transient correlation or time delay (possibly indicating the presence of a regulator that is expressed before the cluster of genes it regulates), as well as inverted profiles (possibly indicating negative regulation).

In addition, many other analyses (both parametric and non-parametric) take only the mean of the replicates into account, effectively ignoring the between-replicate variation. It has been shown the inclusion of the replicate information can greatly improve the analyses (Cooke, et al., 2011). A regression approach is able to use this information as an indicator of the quality of the fitted model in the form of deviations of the functions to the data, as well as the level of uncertainty in the parameter estimates.

In this paper, several methods were developed to fit a variety of models to time-series gene expression data, and then perform a cluster analysis using the fitted parameters. These fitted curves represent a smoothed description of the gene expression profiles from a time-series experiment, and the fitted parameters thus reflect aspects of the underlying biology. By performing a cluster analysis on aspects of the shapes, such as the gradient or time of activation, it is possible to obtain more information regarding the biological processes that are occurring at a given point in time, as compared to using conventional clustering methods. For example, it is possible to identify genes that may be co-regulated by a common transcription factor, or are part of similar metabolic functions. This methodology presents a different philosophy and analysis approach for investigating gene expression profiles, where the profiles are grouped based on important aspects of the profile, instead of simply looking at the entire profile all at once.

The models used were chosen to be representative of the observed gene expression profiles, as well as being intuitive in terms of gene expression processes. For example, the increasing logistic function represents a symmetric sigmoid curve characterised by a rapid growth rate in the beginning, slowing down to a constant growth rate, before finally approaching the asymptotic maximum value.

This analysis was performed on an experiment investigating the gene expression changes occurring in a plant as it undergoes senescence. The results obtained from the analyses suggest that while the selected shapes could adequately describe many of the gene expression profiles, there were still some that were not. Further investigation would be needed to identify and parameterise the missing model shapes. Here, techniques using splines have the advantage, as they are more flexible and thus able to handle unusual profile shapes. However, the purpose of this analysis was to obtain more information from the expression profiles than merely their shapes, namely additional information regarding the underlying mechanisms for the given expression profile. Through the use of the fitted parameters and goodness-of-fit statistics, a more exploratory approach was developed to aid in the analysis of the data.

The thresholds described above were used to identify genes that had a good fit to the data, and were selected based on the number of genes that passed a given threshold. This was in an attempt to maximise the number of genes included in further analyses, while still maintaining a level of stringency. However, these thresholds are still ultimately arbitrary, and may be raised or lowered to make them more or less stringent, respectively.

The cluster analysis, named ShapeCluster, was developed as an application of the fitted models. Using this analysis, it was possible to cluster on aspects of the shape of the expression profiles using different combinations of parameters. This added flexibility to the analysis and allowed for the investigation of the data in multiple ways. Specifically, performing the cluster analysis on an individual parameter permitted the identification of genes that are co-regulated, or participate in response to the biological stress in question.

ShapeCluster operates in a two-part process. First, one of the regression models is selected, and gene expression profiles which fitted this model are used in the second step, namely cluster on one or more of the fitted model parameters. When multiple parameters are used, a number of options are available. In the simplest case, the differences between multiple parameters are combined into a single distance matrix. However, this could result in a loss of the some of the individual underlying structure as described by each parameter. As an alternative, two meta-clustering approaches were devised that would re-cluster the members of each initial cluster using additional information. In so doing, the parameter structure is preserved thus aiding in more effectively identifying co-expressed and potentially co-regulated sets of genes.

The meta-clustering approaches are ideally used with two parameters, although they can be expanded to use more. However, this could lead to clusters with few members. It is possible to combine all the parameters using the simultaneous parameter method, although this would make the clustering more like “traditional” clustering approaches, where the clustering is performed over the entire expression profile, instead of investigating only a specific aspect of the profile, possibly leading to less biologically relevant genes being clustered together.

By using ShapeCluster, it was possible to analyse the data in a number of different ways to identify the biological events that are taking place at a given time. While these methods provide flexibility, it can be over whelming not knowing which shape and parameter combinations should be used. To this end, a table of suggested uses is provided in Table 2.

**Table 2.** Table of recommended shape and parameter combinations in order to investigate a specific biological question.

| Task | Shape | Parameter |
| --- | --- | --- |
| Identify co-regulated genes | Gompertz, logistic | *grad* |
|  | Exponential | *r, b* |
|  | Linear | *m* |
| Identify gene responses with the same response duration | Gaussian | *s* |
| Determine time of activation or repression | Gompertz, logistic | *5per, m* |
|  | Gaussian | *s* |
|  | Linear+exponential | *linpnt* |
|  | Critical exponential | *turnp* |

A slight drawback in this approach is that the number of time points needs to be greater than the number of model parameters (in general, greater than four). Therefore this can be prohibitive for shorter time-series experiments. In addition, this approach was created with microarray data in mind, and not well suited for RNA-Seq count data. However, it may be possible to use data that has been transformed, for example using a variance stabilising transformation.

These analyses provided biologically oriented descriptions of individual gene expression profiles, allowing for the modelling and greater interpretation of profiles obtained from time-series experiments. Through careful choice of appropriate models, such statistical regression approaches allow for an improved comparison of gene expression profiles, and may provide an improved understanding of common regulatory mechanisms between genes. The development of these new tools may provide a better assessment of the mechanisms underlying gene expression responses, and could assist in identifying key time points for further investigation or experiments.

## ACKNOWLEDGEMENTS

Thank you to Sascha Ott and Laura Baxter for assistance with the hypergeometric test analysis.

## REFERENCES

Breeze, E., et al. High-resolution temporal profiling of transcripts during Arabidopsis leaf senescence reveals a distinct chronology of processes and regulation. Plant Cell 2011;23(3):873–894.

Brock, G., et al. clValid, an R package for cluster validation. J Stat Softw 2008;25:i4.

Buchanan-Wollaston, V., et al. Comparative transcriptome analysis reveals significant differences in gene expression and signalling pathways between developmental and dark/starvation-induced senescence in Arabidopsis. Plant J 2005;42(4):567–585.

Burnham, K.P. and Anderson, D.R. Model selection and multimodel inference: a practical information-theoretic approach. New York: Springer; 2002.

Cooke, E.J., et al. Bayesian hierarchical clustering for microarray time series data with replicates and outlier measurements. BMC Bioinformatics 2011;12:399.

Dalbiès-Dulout, A. and Doré, T. Management of inflorescence and viable seed production of blackgrass ( Alopecurus myosuroides) on set-aside in France. Crop Prot 2001;20(3):221–227.

Datta, S. and Datta, S. Methods for evaluating clustering algorithms for gene expression data using a reference set of functional classes. BMC Bioinformatics 2006;7:397.

Déjean, S., et al. Clustering time-series gene expression data using smoothing spline derivatives. EURASIP J Bioinform Syst Biol 2007;2007:70561.

Eastwood, D.C., et al. Statistical modelling of transcript profiles of differentially regulated genes. BMC Mol Biol 2008;9:66.

Falcon, S. and Gentleman, R. Using GOstats to test gene lists for GO term association. Bioinformatics 2007;23(2):257–258. Fox, J. An R and S-Plus Companion to Applied Regression. London; 2002.

Fox, J. Applied Regression Analysis and Generalized Linear Models. SAGE Publications; 2008.

Heard, N., Holmes, C. and Stephens, D. A quantitative study of gene regulation involved in the immune response of anopheline mosquitoes: An application of Bayesian hierarchical clustering of curves. J Am Stat Assoc 2006;101(473):18–29.

Hickman, R., et al. A local regulatory network around three NAC transcription factors in stress responses and senescence in Arabidopsis leaves. Plant J 2013;75(1):26–39.

Higo, K., et al. Plant cis-acting regulatory DNA elements (PLACE) database: 1999. Nucleic Acids Res 1999;27(1):297–300.

Kiddle, S.J., et al. Temporal clustering by affinity propagation reveals transcriptional modules in Arabidopsis thaliana. Bioinformatics 2010;26(3):355–362.

Langfelder, P., Zhang, B. and Horvath, S. Defining clusters from a hierarchical cluster tree: the Dynamic Tree Cut package for R. Bioinformatics 2008;24(5):719–720.

Li, J., et al. A membrane-tethered transcription factor ANAC089 negatively regulates floral initiation in Arabidopsis thaliana. Science China Life Sciences 2010;53(11):1299–1306.

Ma, P., et al. A data-driven clustering method for time course gene expression data. Nucleic Acids Res 2006;34(4):1261–1269.

Mathelier, A., et al. JASPAR 2014: an extensively expanded and updated open-access database of transcription factor binding profiles. Nucleic Acids Res 2014;42(Database issue):D142–147.

Oh, S., et al. Time Series Expression Analyses Using RNA-seq: A Statistical Approach. Biomed Res Int 2013;2013:203681.

Osborne, D.J. Control of leaf senescence by auxins. Nature 1959;183:1459–1460.

Peng, L. and Shikanai, T. Supercomplex formation with photosystem I is required for the stabilization of the chloroplast NADH dehydrogenase-like complex in Arabidopsis. Plant Physiol 2011;155(4):1629–1639.

Ritz, C. and Streibig, J.C. Nonlinear regression with R. New York: Springer; 2008.

Ross, G.J.S. The Efficient Use of Function Minimization in Non-Linear Maximum-Likelihood Estimation. Journal of the Royal Statistical Society 1970;19(3):205–221.

Schliep, A., Schönhuth, A. and Steinhoff, C. Using hidden Markov models to analyze gene expression time course data. Bioinformatics 2003;19 Suppl 1:i255–263.

Schoolfield, R., Sharpe, P. and Magnuson, C. Non-linear regression of biological temperature-dependent rate models based on absolute reaction-rate theory. J Theor Biol 1981;88(4):719–731.

Thimm, O., et al. mapman: a user-driven tool to display genomics data sets onto diagrams of metabolic pathways and other biological processes. Plant J 2004;37(6):914–939.

van der Graaff, E., et al. Transcription analysis of Arabidopsis membrane transporters and hormone pathways during developmental and induced leaf senescence. Plant Physiol 2006;141(2):776–792.

Watkins, P. and Venables, W.N. Non-linear regression for optimising the separation of carboxylic acids. R News 2006;6:2–7.

Woolcock, A.J., Salome, C.M. and Yan, K. The shape of the dose-response curve to histamine in asthmatic and normal subjects. Am Rev Respir Dis 1984;130(1):71–75.

